# Anti-CRISPR AcrIIC5 is a dsDNA mimic that inhibits type II-C Cas9 effectors by blocking PAM recognition

**DOI:** 10.1101/2023.01.17.524369

**Authors:** Wei Sun, Xiaolong Zhao, Jinlong Wang, Xiaoqi Yang, Zhi Cheng, Shuo Liu, Jiuyu Wang, Gang Sheng, Yanli Wang

## Abstract

Anti-CRISPR proteins are encoded by phages to inhibit the CRISPR-Cas systems of the hosts. AcrIIC5 inhibits several naturally high-fidelity type II-C Cas9 enzymes, including orthologs from *Neisseria meningitidis* (Nme1Cas9) and *Simonsiella muelleri* (SmuCas9). Here, we solve the structure of AcrIIC5 in complex with Nme1Cas9 and sgRNA. We show that AcrIIC5 adopts a novel fold to mimic the size and charge distribution of double-stranded DNA, and uses its negatively charged grooves to bind and occlude the protospacer adjacent motif (PAM) binding site in the target DNA cleft of Cas9. AcrIIC5 is positioned into the crevice between the WED and PI domains of Cas9, and one end of the anti-CRISPR interacts with the phosphate lock loop and a linker between the RuvC and BH structural domains. We employ biochemical and mutational analyses to build a model for AcrIIC5’s mechanism of action, and identify residues on both the anti-CRISPR and Cas9 that are important for their interaction and inhibition. Together, the structure and mechanism of AcrIIC5 reveal convergent evolution among disparate anti-CRISPR proteins that use a DNA-mimic strategy to inhibit diverse CRISPR-Cas surveillance complexes, and provide new insights into a tool for potent inhibition of type II-C Cas9 orthologs.

## Introduction

CRISPR-Cas systems are adaptive immune systems of bacteria and archaea that defend against bacterial viruses (bacteriophages) and other mobile genetic elements (MGEs) (1-3). CRISPR-Cas systems capture DNA fragments from invading DNA and integrate them into the CRISPR array as new spacers. CRISPR arrays are transcribed and processed into mature CRISPR RNAs (crRNAs) (4,5) that guide the surveillance complex of CRISPR-associated (Cas) proteins to cleave foreign nucleic acids (6-8). CRISPR-Cas systems are categorized into two classes and six types based on phylogeny, the set of *cas* genes present, and their mechanisms of action (9). Class I (types I, III, and IV) includes systems with multi-subunit surveillance, whereas Class II systems (types II, V, and VI) consist of a single effector protein, including the well-studied type II Cas9 effectors.

To evade the threat of CRISPR-Cas systems in their host bacteria, some bacteriophages and other MGEs encode anti-CRISPR (Acr) proteins (10-12). The ‘arms race’ between phages and bacteria over millions of years of evolution has given rise to Acrs against nearly all types of CRISPR-Cas systems, with likely many more awaiting discoveries (13-20). Acrs employ various mechanisms to inhibit CRISPR-Cas systems, such as hindering crRNA loading (21,22), preventing target DNA or RNA binding (20,23), inhibiting the active site of the nuclease domain (24), or blocking the conformational changes required to activate the cleavage-competent state of the surveillance complex (25,26).

Despite the widespread and successful adaptation of Cas9- and other CRISPR-based tools for gene editing (27,28), the significant drawback of unwanted off-target cleavage remains a challenge. Many high-fidelity variants of Cas proteins have been engineered to address this issue, and characterization of new orthologs from bacterial systems has yielded some promising new candidates with naturally high fidelity, including many enzymes in the type II-C clade of the Cas9 tree (29). Among the known type II-C Cas9 orthologs, two paralogs from *Neisseria meningitidis*, Nme1Cas9 and Nme2Cas9, have been validated for genome engineering in human cells (30-32).

Acrs provide an alternative approach to control CRISPR-Cas activity, and therefore hold promise to reduce off-target editing resulting from excessive or prolonged Cas protein activity in cells (33-35). Moreover, Acr inhibition mechanisms can help us better understand the conformational changes and structural stages of CRISPR-Cas activation by trapping effectors in different states and providing reagents to specifically block particular biochemical activities of Cas proteins.

Focusing our efforts here on the high-fidelity type II-C Cas9 enzymes, there are five known Acrs that inhibit this subtype (AcrIIC1-AcrIIC5) and the mechanisms of four of these five have been elucidated (14,16). AcrIIC1 binds to the catalytic sites in the HNH nuclease domain and masks the RuvC domain to prevent target DNA cleavage (24). AcrIIC2 binds to the positively-charged BH domain and prevents guide RNA loading, thereby blocking surveillance complex assembly (21,22). Two AcrIIC3 proteins tether two Cas9 proteins together by interacting with the HNH and REC2 domains (26), thereby reducing the mobility of the HNH domain and thus preventing Cas9 activation. AcrIIC4, on the other hand, prevents the recruitment of target DNA to Cas9 (16,36). AcrIIC5 was reported to be the most potent type II-C Cas9 inhibitor, but its mechanism of action and its structure remain elusive. This Acr is particularly interesting since it can inhibit the type II-C effectors from *Simonsiella muelleri* (SmuCas9), *Haemophilus parainfluenzae* (HpaCas9), and Nme1Cas9, but not the closely related Nme2Cas9 (16,32).

In this study, we determined the cryogenic electron microscopy (cryo-EM) structure of AcrIIC5 bound to the Nme1Cas9:sgRNA complex and performed mutational and biochemical analyses to determine its mechanism of action. We find that AcrIIC5 only binds to the sgRNA-bound Nme1Cas9 surveillance complex and not apo-Nme1Cas9. AcrIIC5 does not resemble any known protein fold, but its globular structure has similar dimensions, shape, and charge distribution as a segment of double-stranded DNA (dsDNA). Our structure reveals that AcrIIC5 binds to the cleft between the WED and PI domains of Cas9, occupying the site where the double-stranded protospacer adjacent motif (PAM) of the target DNA would bind. The inability of AcrIIC5 to inhibit the Nme2Cas9 paralog can be explained by a subtle change in the sequence of the phosphate lock loop of Nme2Cas9 compared to Nme1Cas9 and SmuCas9. Our results define the structure and mechanism of a potent anti-CRISPR dsDNA mimic, and suggest that evasion of Acr binding may drive the evolution of Cas proteins.

## Materials and methods

### Protein expression and purification

Full-length ORFs of Nme1Cas9 (encoding residues 1-1082), Nme2Cas9 (encoding residues 1-1082), SmuCas9 (encoding residues 1-1065) and AcrIIC5 (encoding residues 1-130) were synthesized from Sangon Biotech and cloned into an expression vector pET28a-Sumo with a N-terminal His_6_-Sumo tag. The codons of these genes are optimized according to *Escherichia coli*. Mutants were constructed using a site-directed mutagenesis kit. All proteins were overexpressed in *E. coli* BL21 (DE3) (Novagen) cells and were induced with 0.2 mM isopropyl-β-D-1-thiogalactopyranoside (IPTG) at OD_600_ = 0.8 at 18°C for 12 h.

Cells expressing Cas9 were harvested and then disrupted by sonication in lysis buffer containing 20 mM Tris-HCl (pH 7.5) and 0.5 M NaCl at 4°C following high-speed centrifugation. The supernatant was incubated with Ni^2+^-Sepharose resin (GE Healthcare), and the bound protein was eluted with lysis buffer supplemented with 200 mM imidazole. The eluted protein was incubated with His-tagged ubiquitin-like protein 1 (Ulp1) protease to remove the His_6_–Sumo-tag, and dialyzed against buffer containing 20 mM Tris-HCl (pH 7.5), 250 mM NaCl at 4°C for 2 h. Cas9 proteins were further purified by chromatography on Ni-NTA and SP column (GE Healthcare).

Cells expressing AcrIIC5 were disrupted by sonication in lysis buffer containing 20 mM Tris-HCl (pH 7.5) and 0.5 M NaCl at 4°C. After high-speed centrifugation, the supernatant was incubated with Ni^2+^-Sepharose resin (GE Healthcare), and the bound protein was eluted with lysis buffer supplemented with 250 mM imidazole. His_6_-Sumo tag was cleaved from AcrIIC5 protein by His-tagged Ulp1 protease during dialysis against buffer containing 20 mM Tris-HCl (pH 7.5), 200 mM NaCl for 2 h at 4°C and removed by a second Ni^2+^-Sepharose column. The flow-through collections were loaded into a Q column (GE Healthcare) equilibrated by buffer containing 20 mM Tris-HCl (pH 7.5), 200 mM NaCl, eluting with buffer containing 20 mM Tris-HCl (pH 7.5), 1 M NaCl.

### In vitro transcription and purification of sgRNA

The 135-nucleotide (nt) sgRNA of NmeCas9 was synthesized by in vitro transcription with T7 RNA polymerase and plasmid DNA templates linearized by HindIII-HF (New England Biolabs). The sgRNA sequence was listed in Supplementary Table S1. Transcription reactions were performed at 37°C for 4 h in buffer containing 100 mM HEPES-KOH (pH 7.9), 30 mM MgCl_2_, 30 mM DTT, 3 mM each NTPs, 2 mM spermidine, 0.05 mg/mL T7 RNA polymerase, and 35 ng/μL linearized plasmid DNA template. The RNA was then separated on a 12% denaturing (8 M urea) polyacrylamide gel and concentrated via an Elutrap system. Finally, RNA was dissolved in DEPC (diethylpyrocarbonate) H_2_O and stored at -40°C. Nme1Cas9, Nme2Cas9 and SmuCas9 use the same 135-nt sgRNA.

### Preparation of DNA

All short DNAs used in this study were purchased from Sangon Biotech. The DNA sequences used for cleavage and binding assays were shown in Supplementary Table S1. All DNAs were dissolved in buffer containing 20 mM Tris-HCl (pH 7.5), 150 mM NaCl, 10 mM MgCl_2_. For target dsDNA, the target strand (TS) and the nontarget strand (NTS) were mixed at a molar ratio of 1:1.05, pre-denatured at 95°C for 10 min and then annealed at room temperature.

### Size Exclusion Chromatography (SEC) binding assays

SEC binding assays were performed using a Superdex 200 increase Hiload 10/300 column (GE healthcare) at a flow rate 0.4 ml/min with absorbances monitored at 280 nm. Nme1Cas9, sgRNA, DNA, and AcrIIC5 were mixed at a molar ratio of 1:∼1.1:1.2:5. All components were added one by one according to the order in which they appear in the complex name and incubated on ice for 30 min before adding next component. All assays were performed in buffer containing 20 mM Tris (pH 7.5), 250 mM NaCl. Fractions from all assays were analyzed on 15% SDS-PAGE stained with Coomassie blue. The nucleic acids of some fractions were subjected to phenol-chloroform extraction and analyzed through 20% denaturing Urea-PAGE visualized by toluidine blue staining. All assays were repeated at least three times.

### In vitro cleavage assay

pUC19 target DNA (35 bp target DNA cloned into a modified pUC19 vector) was used to test the dsDNA cleavage activity of various Cas9 orthologs and the inhibition potency of AcrIIC5. The modified pUC19 target DNA plasmid was linearized by HindIII-HF (New England Biolabs) in advance. Prepared complexes Cas9-sgRNA or Cas9-sgRNA-AcrIIC5 were incubated with 300 ng pUC19 target DNA in 10 μL reaction buffer containing 20 mM Tris-HCl (pH 7.5), 100 mM KCl, 10 mM MgCl_2_, 1 mM DTT and 5% glycerol. The ratio of Cas9:sgRNA was 1:1.1, and the final concentration of Cas9 was 100 nM unless illustrated individually. Reactions were incubated at 37°C for 5-60 min and were quenched by 2 μL 6× loading dye containing 10 mM EDTA. Products were loaded into 1% agarose gels. DNA was stained with ethidium bromide and visualized by a UV spectrometer. All experiments were repeated at least three times.

### Electrophoretic mobility shift assay (EMSA)

First, 3 μL 10 μM Nme1Cas9 (D16A/H588A) was mixed with 3 μL 10 μM sgRNA in 1× binding buffer [20 mM Tris (pH7.5), 250 mM NaCl] and incubated at 25°C for 20 min to reconstitute the binary complex. Subsequently, 3 μL AcrIIC5 of various concentrations (0, 5, 10, 20, 50, 100, 200 μM) was added for another 20 min incubation at 25°C. Then, 3 μL 2 μM Cy3-labeled dsDNA was added for 20 min incubation at 37°C. Finally, 2.4 μL 6× loading buffer [50 mM Tris (pH 8.5), 500 mM glycine, 48% glycerol] was added to the sample. 10 μL each sample was then loaded onto a 5% native polyacrylamide gel, and the Cy3-labeled dsDNA was visualized using a FluorChem system (Protiensimple). The experiment was repeated at least three times.

### Reconstitution of Nme1Cas9-sgRNA-AcrIIC5 complex

The Nme1Cas9-sgRNA-AcrIIC5 complex was reconstituted by incubating Nme1Cas9 with sgRNA on ice for 30 min, and then AcrIIC5 was added for incubation for another 30 min. The molar ratios of Nme1Cas9, sgRNA and AcrIIC5 were 1:1.1:5. Next, the resulting sample was loaded into a Superdex 200 increase Hiload 10/300 column (GE Healthcare) in buffer containing 20 mM Tris (pH 7.5), 150 mM NaCl. Finally, the complex was concentrated to an A_280_ absorbance of 0.8, as measured by Nanodrop 2000, prior to cryo-EM grid preparation.

### Cryo-EM sample preparation and data acquisition

C411 Cu 400 mesh grids were glow discharged in O_2_-Ar condition for 60 s. 3 μL solution of Nme1Cas9-sgRNA-AcrIIC5 was applied to the grid followed by blotting for 6.0 s at 100% humidity and 4 °C, and flash-frozen in liquid ethane using a Vitrobot Mark IV (Thermo Fisher Scientific, USA).

Grids were imaged with a 300 kV Titan Krios (Thermo Fisher Scientific, USA) equipped with a K2 Summit direct electron detector (Gatan, USA) and a GIF-Quantum energy filter. Dose-fractionated super-resolution movie stacks collected with a calibrated magnification of 130,000× were binned to a pixel size of 1.04 Å. The defocus range was set to between -1.0 and -1.6 μm. Each movie stack was collected using SerialEM with a total exposure time of 6.4 s, and dose-fractioned to 32 frames, resulting in a total dose of 60 e-/Å^2^.

### Single-particle cryo-EM data processing

A total of 3197 movie stacks were collected and all image processing was performed in cryoSPARC v3.1 (37). Patch motion correction, CTF-estimation, template picking, 2D classification, 3D classification and non-uniform 3D refinement were performed in chronological order. A total of 2,685,237 particles were automatically picked by template picker and extracted with a 260^2^-pixel box in cryoSPARC. After 3 rounds of reference-free 2D classification, 291,265 particles were selected for Ab-Initio reconstruction and heterogeneous refinement. Finally, all the 291,265 particles were used to generate the final map at a resolution of 3.09 Å reported according to the golden-standard Fourier shell correlation (GSFSC) criterion.

### Model fitting and refinement

The atom model of the complex was generated by first fitting the chains of Nme1Cas9-sgRNA binary complex (PDB: 6JDQ) and structure of apo-AcrIIC5 predicted by AlphaFold2 (38) into the cryo-EM density in Chimera (39). Then, the atom model was manually adjusted and corrected according to the protein sequences and cryo-EM densities in Coot (40), and subsequently, real-space refinement was performed by PHENIX (41). Details of the data collection and refinement statistics of the complex are summarized in Supplementary Table S2.

All structure figures were prepared using PyMOL (42) or UCSF ChimeraX (43).

## Results

### AcrIIC5 binds to Nme1Cas9-sgRNA and prevents the binding of dsDNA to inhibit cleavage

AcrIIC5, which was originally identified from a prophage of *Simonsiella muelleri*, is an anti-CRISPR protein that counteracts the type II-C CRISPR-Cas system (16). Our in vitro DNA cleavage assay in the presence of AcrIIC5_Smu_ (referred to as AcrIIC5 hereafter) shows that AcrIIC5 exhibits cross-species inhibitory activity (Figure 1A). We found that AcrIIC5 was able to inhibit the activity of Nme1Cas9 and SmuCas9, which possess 5’-N_4_GATT and 5’-N_4_C PAM preferences, respectively. In contrast, AcrIIC5 fails to show any inhibitory activity against Nme2Cas9 that possesses 5’-N_4_CC PAM preference, despite its high degree of sequence similarity with Nme1Cas9 throughout most of the protein apart from the C-terminal WED and PI domains. These observations suggest that the protein sequence within the C-terminal region of Cas9 may be critical for AcrIIC5.

**Figure 1.**
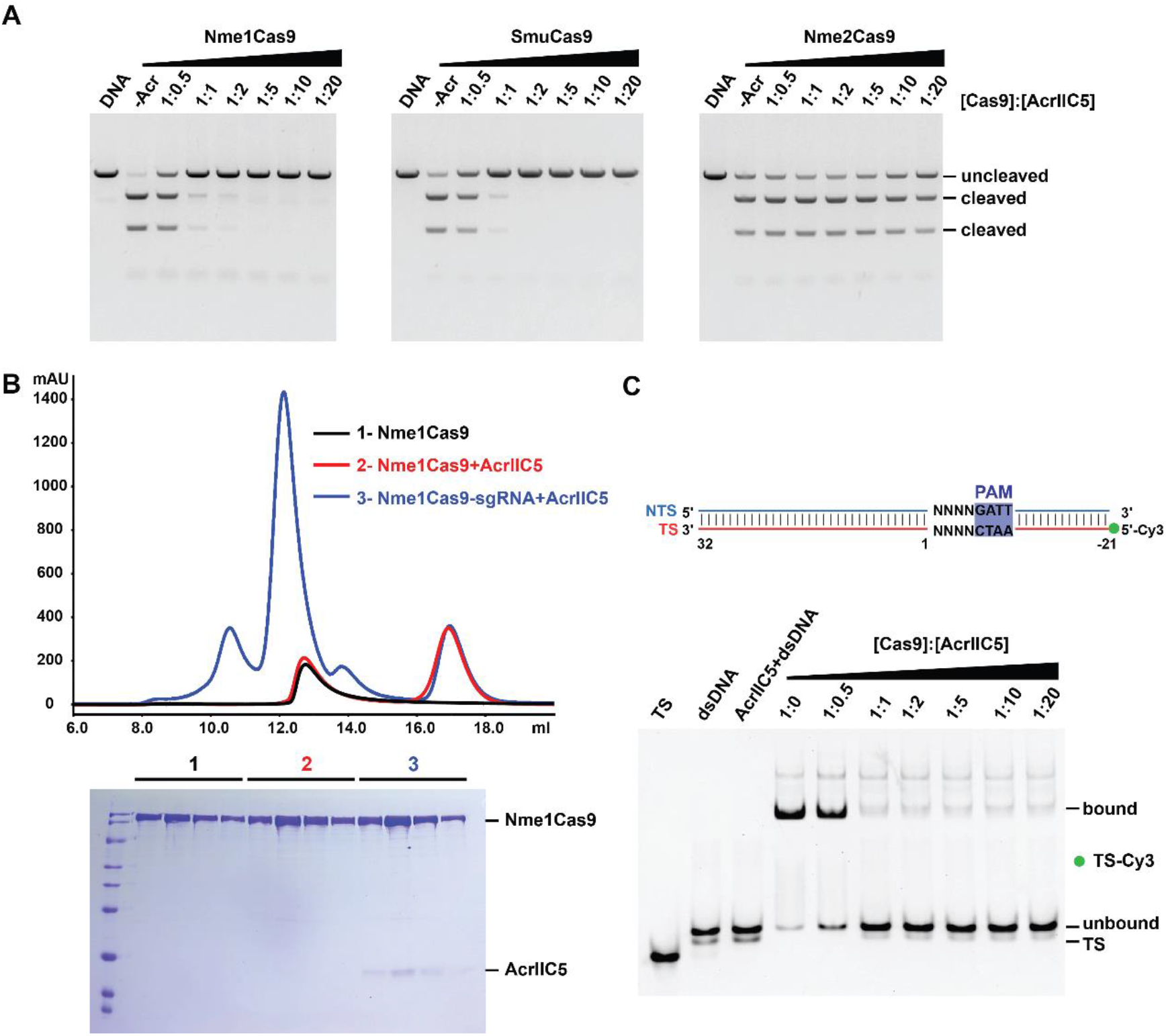
AcrIIC5 binds the Nme1Cas9-sgRNA complex to inhibit dsDNA cleavage. (A) In vitro DNA cleavage assay of Nme1Cas9, SmuCas9, and Nme2Cas9, without or with varying concentrations of AcrIIC5, using a linearized plasmid substrate. The PAM preferences of Nme1Cas9, SmuCas9, and Nme2Cas9 are 5’-N_4_GATT, 5’-N_4_C and 5’-N_4_CC, respectively. The molar ratios of Cas9 to AcrIIC5 were 1:0.5, 1:1, 1:2, 1:5, 1:10 and 1:20. Cleavage was performed at 37°C for 10 min and evaluated using 1% agarose gel stained with ethidium bromide. The figure is a representative of three replicates. (B) Size-exclusion chromatography curves of apo-Nme1Cas9 (black), Nme1Cas9+AcrIIC5 (red), and Nme1Cas9-sgRNA+AcrIIC5 (blue) were overlaid. Fractions from the Cas9-containing peaks at volume of 11-13 mL were characterized by SDS-PAGE and stained with Coomassie blue. The gel is placed at the bottom of the curves. (C) Electrophoretic mobility shift assay to assess the binding of target dsDNA to the Nme1Cas9-sgRNA complex in the presence of AcrIIC5 of various concentrations. A schematic of the 53-bp dsDNA with TS labeled with 5’-Cy3 used in this assay is shown on the top. AcrIIC5+dsDNA is the AcrIIC5 mixed with dsDNA directly, with no Cas9:sgRNA present, as a negative control. The molar ratios of Cas9 to AcrIIC5 are shown above the gel. The figure is a representative of three replicates.

To gain insight into how AcrIIC5 binds to Cas9, we performed size exclusion chromatography (SEC) of a mixture of AcrIIC5 and Nme1Cas9 or SmuCas9 in either the sgRNA-bound or apo state. We found that AcrIIC5 binds sgRNA-bound Cas9 but not apo-Cas9 of both orthologs (Figure 1B and Supplementary Figure S1), mirroring previous results with two anti-CRISPRs against the type II-A CRISPR-Cas system, AcrIIA2 and AcrIIA4 (23,44-46). This result indicates that the AcrIIC5 affects the target DNA binding or activation of Cas9.

Next, to test whether AcrIIC5 prevents target dsDNA binding to the Nme1Cas9-sgRNA complex, we used in vitro competition assays to investigate AcrIIC5 and target DNA binding to Nme1Cas9 (Supplementary Figure S2). Due to the low binding affinity of fully-complementary dsDNA with Nme1Cas9, partially-duplexed dsDNA was used for this competition binding assay (Supplementary Figure S2A). When DNA was mixed with the dead Nme1Cas9-sgRNA complex first before AcrIIC5 addition, there was no obvious binding between AcrIIC5 and Nme1Cas9. By contrast, when AcrIIC5 was pre-bound to Nme1Cas9-sgRNA complex, the target DNA failed to bind to Nme1Cas9-sgRNA complex (Supplementary Figure S2B-S2D). We next performed an electrophoretic mobility shift assay (EMSA) by adding a constant amount of Cy3-labeled fully complementary dsDNA to the pre-formed Cas9-sgRNA complex in the presence of different concentrations of AcrIIC5. We saw a dose-dependent effect of increasing concentrations of AcrIIC5 on the inhibition of dsDNA binding to Cas9, with no detectable binding when the molar ratio of Cas9 to AcrIIC5 reached 1:1 (Figure 1C). Together, these data suggest that AcrIIC5 competes with dsDNA for the binding site in the Nme1Cas9-sgRNA effector complex or prevents dsDNA binding allosterically.

### Cryo-electron microscopy structure of Nme1Cas9-sgRNA-AcrIIC5 ternary complex

To elucidate the molecular mechanism of the inhibition of Nme1Cas9 by AcrIIC5, we solved the structure of the Nme1Cas9-sgRNA-AcrIIC5 ternary complex using single-particle cryo-electron microscopy (cryo-EM) at a resolution of 3.09 Å (Supplementary Figure S3). Nme1Cas9 is a multi-domain protein composed of 1082 amino acids and is divided into the REC and NUC lobes (Figure 2A). A 135-nt single-guide RNA (sgRNA) was used for complex reconstitution (Figure 2B), and we confirmed that the sgRNA in our structure is assembled with Nme1Cas9 in a similar manner as previously observed in the Nme1Cas9-sgRNA binary complex (PDB: 6JDQ) (26). Only 9-nt of the spacer was observed in our structure, while the other 15 nucleotides could not be exactly modeled. The Nme1Cas9-sgRNA-AcrIIC5 complex revealed one AcrIIC5 monomer bound to one NmeCas9-sgRNA binary complex (Figure 2C). The structural comparison between Nme1Cas9-sgRNA-AcrIIC5 and Nme1Cas9-sgRNA complexes does not indicate major conformational changes of Nme1Cas9 upon binding to AcrIIC5 (Figure 2D).

**Figure 2.**
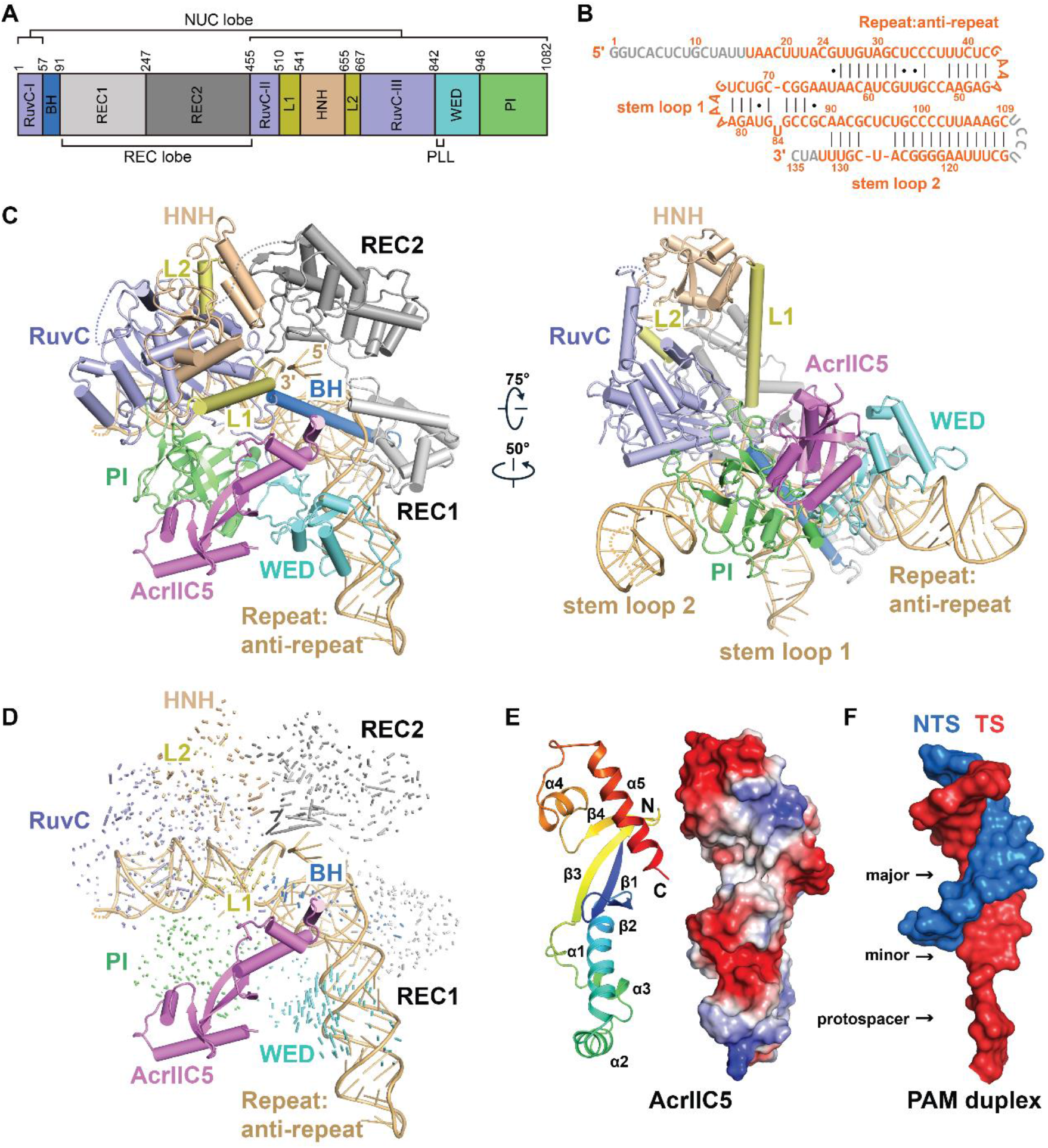
Cryo-EM structure of Nme1Cas9-sgRNA-AcrIIC5 ternary complex. (A) Domain organization of Nme1Cas9. (B) Schematic of the sgRNA used for complex reconstitution. Nucleotides that cannot be traced in the electron density map are colored in gray. (C) Overall structure of Nme1Cas9-sgRNA-AcrIIC5 complex. Individual Nme1Cas9 domains are colored according to the scheme in (A). AcrIIC5 is colored in violet. Two separate views are displayed. (D) Structural comparison between Nme1Cas9-sgRNA-AcrIIC5 and Nme1Cas9-sgRNA (PDB: 6JDQ) complexes. (E) Cartoon and surface representations of AcrIIC5 in the structure of Nme1Cas9-sgRNA-AcrIIC5. (F) Surface representation of the PAM duplex from the structure of Nme1Cas9 H588A-sgRNA-dsDNA (PDB: 6KC7). For the PAM duplex, the 11 base-pair (bp) TS:NTS duplex and 3 nucleotide (nt) protospacer moiety of the TS after unwinding are shown.

AcrIIC5 consists of 130 amino acids and is composed of four β-strands and five α-helices. The four-stranded β-sheet is located in the center and flanked by α1-3 on one side and α4-5 on the other. No similar structure was found through a search by the DALI server, indicating that AcrIIC5 has a novel fold. The overall shape of AcrIIC5 structure resembles a long, narrow bundle of sticks with two concave grooves on its surface (Figure 2E). AcrIIC5 is an acidic protein with theoretical pI 4.39 and its two grooves are negatively charged. Interestingly, these two grooves on the surface of AcrIIC5 resemble the shapes of the major and minor grooves of the PAM duplex, which resembles the conventional double-stranded DNA duplex. We refer to these as the major and minor concave surfaces of AcrIIC5 from here forward (Figure 2F).

### AcrIIC5 is positioned in the cleft between the WED and PI domains

In the cryo-EM structure of the Nme1Cas9-sgRNA-AcrIIC5 complex, AcrIIC5 is located in the crevice between the WED and PI domains, where the middle region of the Acr forms extensive interactions with both domains (Figure 3A). The PI domain is embedded in the major concave surface of AcrIIC5 and contacts AcrIIC5 primarily through charge-charge interactions. Specifically, residues K1012, K1013, and K1043 within the PI domain interact with residues E100, D97, and E26 of AcrIIC5, respectively (Figure 3B). The WED domain contacts the minor concave surface of AcrIIC5, with Cas9 residues N944 and D943 forming hydrogen bonds or salt bridges with Y11 and R16 of AcrIIC5, respectively (Figure 3C). In addition, the helix-turn-helix (amino acids 22-52) of AcrIIC5 inserts into the crevice formed by the WED, RuvC and BH domains, primarily contacting the phosphate lock loop (PLL) within the WED domain – which is critical for target dsDNA unwinding and initial base-paring with the seed region of the sgRNA (Supplementary Figure S4) (47-49) and the linker connecting the RuvC nuclease and BH domains (RuvC-BH linker) (Figure 3D). Specifically, the side-chain of D36 in AcrIIC5 forms three hydrogen bonds with residues E845 and T846 of the PLL. In addition, the side-chains of E845 and T846 form hydrogen bonds with the main-chain of AcrIIC5 residues G40 and Y41, and the side-chain of N39, respectively. Moreover, AcrIIC5 residues E37, K49, and S69 contact with residues K53, D56, and P52 within the RuvC-BH linker, respectively.

**Figure 3.**
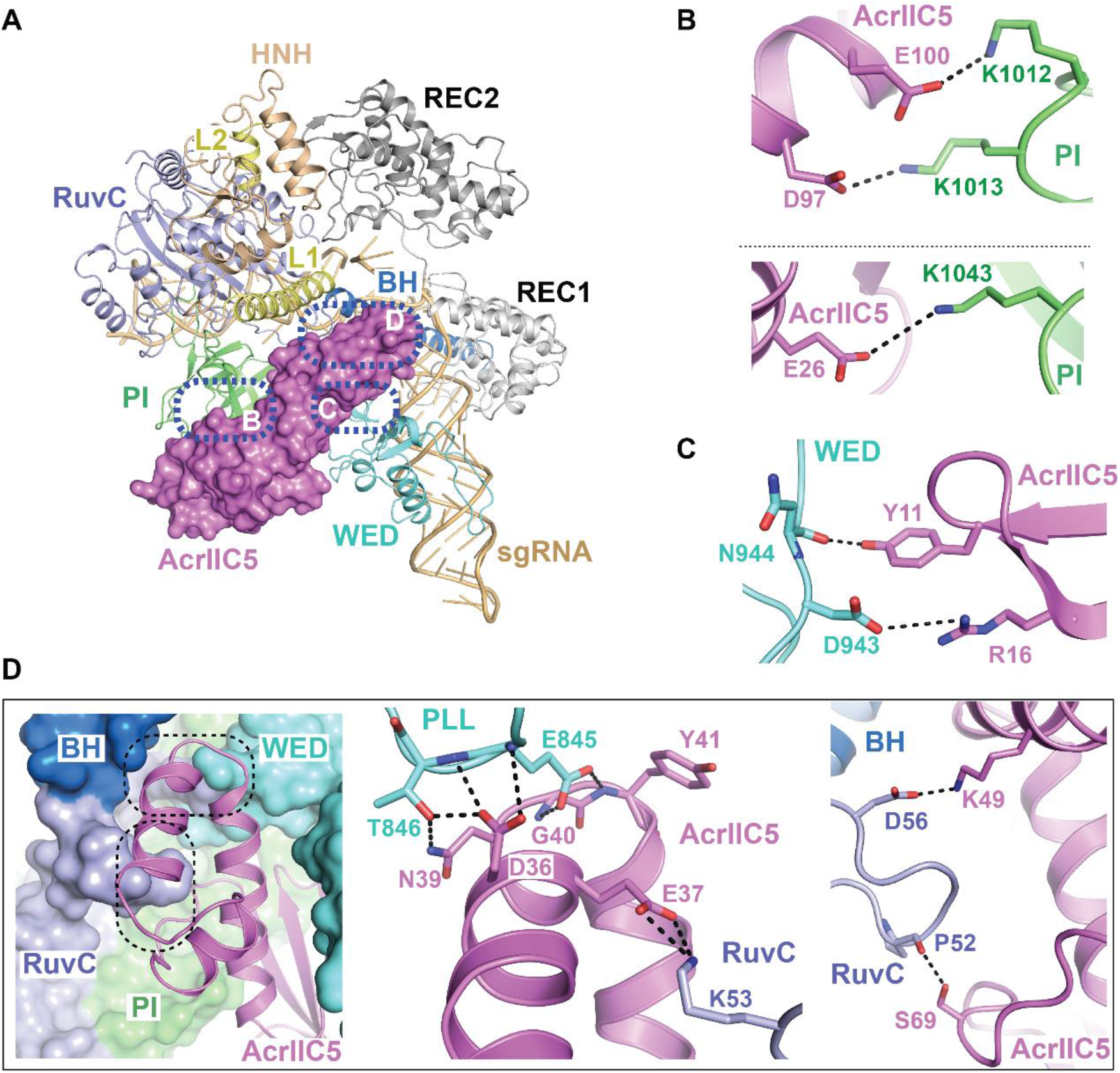
Interactions between AcrIIC5 and Nme1Cas9. (A) Overall view of the structure of Nme1Cas9-sgRNA-AcrIIC5 complex. Nme1Cas9 and sgRNA are displayed as cartoon and AcrIIC5 as surface. Interaction surfaces between AcrIIC5 and Nme1Cas9 are highlighted with blue dashed-line circles. (B) AcrIIC5 interacts with the PI domain of Nme1Cas9 through salt bridges. The key residues involved in the interaction are shown as sticks. (C) Interactions between AcrIIC5 and the WED domain of Nme1Cas9. (D) AcrIIC5 inserts into the crevice established by the PLL and RuvC-BH linker. AcrIIC5 is shown as cartoon and Nme1Cas9 is shown as a surface representation. The details of two interfaces are further displayed in two close-up views.

To characterize the importance of key residues of AcrIIC5 involved in binding to Nme1Cas9, we built a panel of alanine substitution mutants of the Acr. We then conducted in vitro DNA cleavage assays in the presence of these AcrIIC5 variants with a molar ratio of 1:10 for Cas9 versus AcrIIC5. We found that single mutation of D36 or E37 impaired the inhibitory activity of AcrIIC5, while the D36A/E37A double mutant completely abolished its activity and resembled the no-Acr control assay (Figure 4A). In contrast, the single point mutations of other tested Cas9-interacting residues in our structure (R16A, E26A, N39A, K49A, S69A, and D97A) displayed no obvious effect when compared to wild-type AcrIIC5. Furthermore, in contrast to wild-type AcrIIC5, the D36A/E37A double mutant failed to bind to the Nme1Cas9-sgRNA complex as assessed by SEC (Figure 4B). In addition, the D36A/E37A double mutant also impaired the inhibitory effect of AcrIIC5 on SmuCas9 (Supplementary Figure S5). Together, these results suggest that residues D36 and E37 of AcrIIC5 are critical for its binding to and inhibition of Nme1Cas9 and SmuCas9.

**Figure 4.**
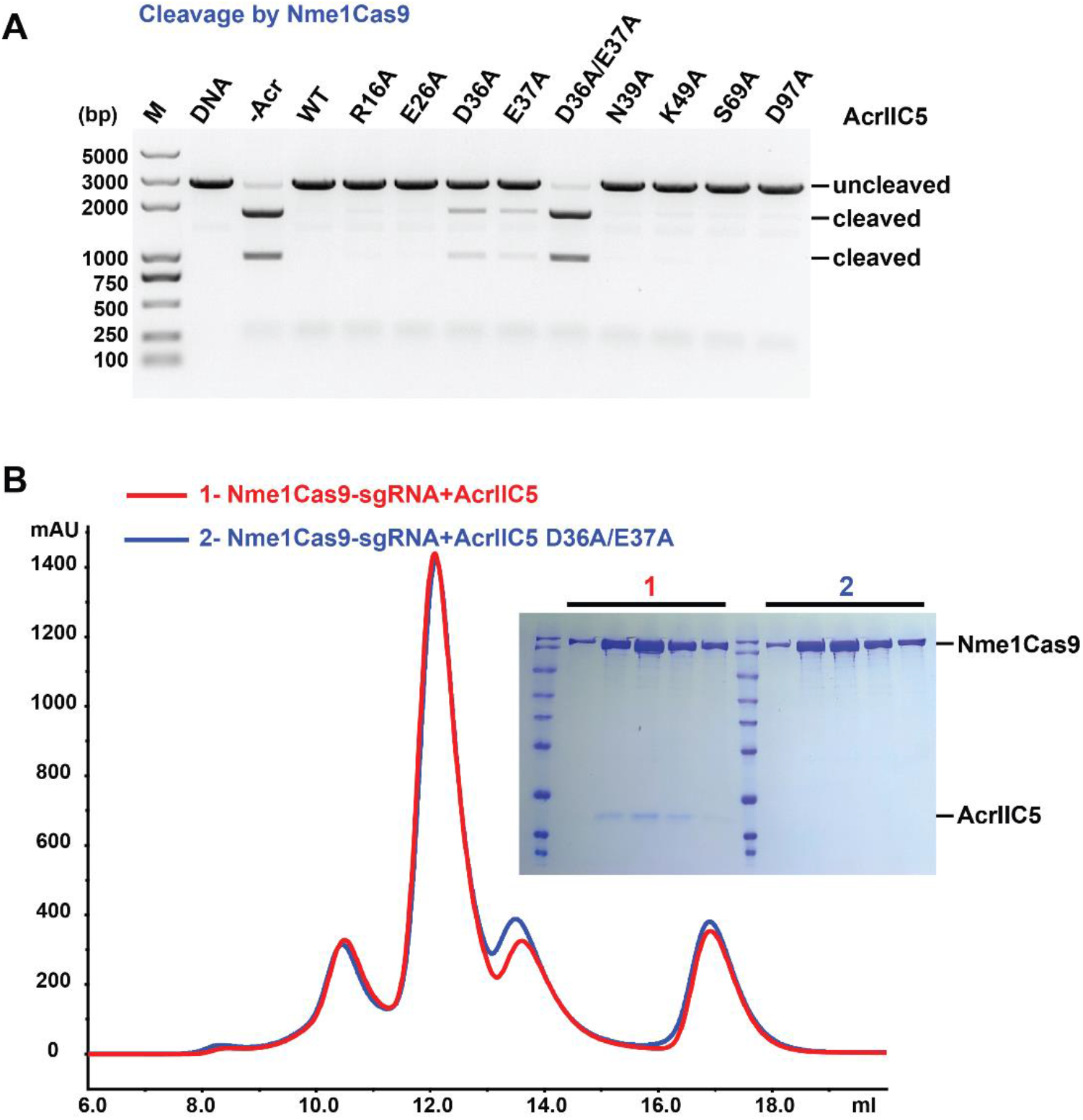
Characterization of key residues of AcrIIC5 involved in binding to Nme1Cas9. (A) In vitro cleavage assay of Nme1Cas9 on a linearized plasmid substrate in the presence of wild-type, single mutants, or double mutants of AcrIIC5. The molar ratio of Cas9 to AcrIIC5 is 1:10. Cleavage was performed at 37°C for 60 min. The resultant DNA fragments were analyzed using 1% agarose gel followed by ethidium bromide staining. The figure is a representative of three replicates. (B) Size-exclusion chromatography curves of Nme1Cas9-sgRNA+AcrIIC5 (red) and Nme1Cas9-sgRNA+AcrIIC5 D36A/E37A (blue) are overlaid. Fractions from the main peaks were characterized by SDS-PAGE followed by Coomassie blue staining.

### AcrIIC5 occupies the PAM binding site

Structural comparison of AcrIIC5-bound and dsDNA-bound Nme1Cas9-sgRNA (PDB: 6KC7) complexes revealed that AcrIIC5 and the PAM duplex moiety of target dsDNA reside in a very similar position of Nme1Cas9 (Figure 5A). The superposition of the two structures showed that AcrIIC5 occupies the position ranging from (+3) to (-11) base pairs of dsDNA (Figure 5B). These findings suggest that AcrIIC5 and dsDNA compete for the same binding position in the Nme1Cas9-sgRNA complex. This is consistent with our SEC binding assay showing that AcrIIC5 and dsDNA compete each other for binding to Nme1Cas9-sgRNA (Supplementary Figure S2).

**Figure 5.**
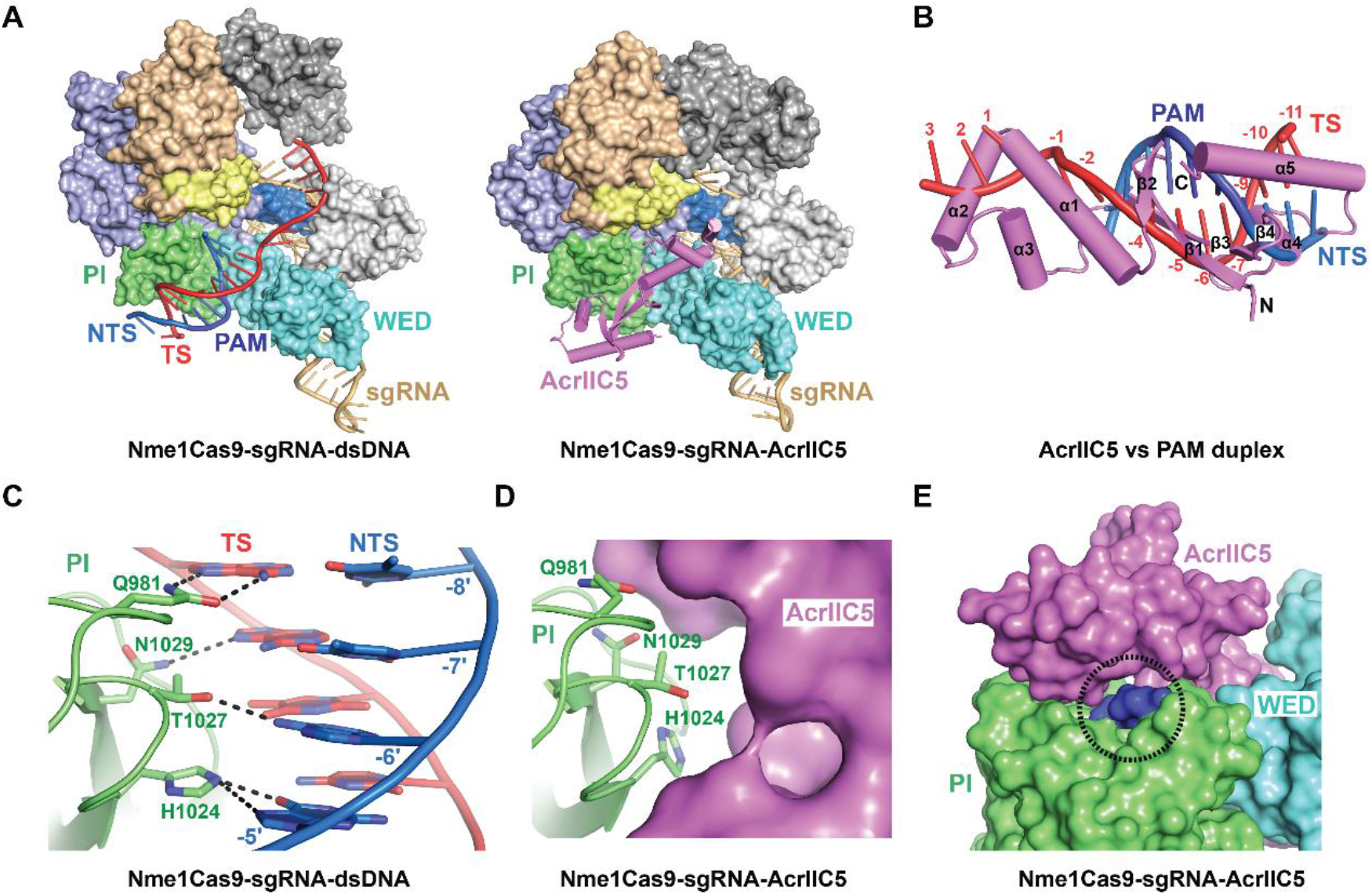
AcrIIC5 prevents binding of dsDNA to Nme1Cas9-sgRNA through hinderance of PAM recognition. (A) Structural comparison between Nme1Cas9 H588A-sgRNA-dsDNA complex (PDB: 6KC7) (left panel) and the Nme1Cas9-sgRNA-AcrIIC5 complex (right panel). (B) Superposition of structures of Nme1Cas9-sgRNA-AcrIIC5 and Nme1Cas9 H588A-sgRNA-dsDNA (PDB: 6KC7) complexes. The PI domains of the two structures were chosen as the reference. Only AcrIIC5 (violet) and the PAM duplex (red and blue) are displayed. (C) Interactions between the PAM sequence and Nme1Cas9 in the structure of Nme1Cas9 H588A-sgRNA-dsDNA (PDB: 6KC7). (D) PAM recognition residues are covered by AcrIIC5 in the structure of Nme1Cas9-sgRNA-AcrIIC5. (E) AcrIIC5 covers the PAM recognition residues in the structure of Nme1Cas9-sgRNA-AcrIIC5. The surface of the PI domain comprised of the PAM recognition residues is shown in blue and highlighted by a dashed-line circle.

Residues Q981, H1024, T1027, and N1029 within the PI domain of Nme1Cas9 are crucial for recognition of the PAM sequence (Figure 5C) (26). Our structure shows that AcrIIC5 binding to the PI domain conceals these PAM recognition residues (Figure 5D), although we did not observe direct interactions between AcrIIC5 and the specific PAM recognition residues, indicating that AcrIIC5 uses different contacts to bind the same surface. As a result, the residues that recognize PAM sequence are no longer solvent exposed, blocking the ability of the PI domain to recognize the PAM – a critical first step in target DNA binding (Figure 5E).

### The phosphate lock loop is essential for the AcrIIC5 inhibition

Next, we asked whether our structural and mutational insights could explain AcrIIC5’s inability to inhibit the second well-characterized *N. meningitidis* Cas9 paralog, Nme2Cas9. To this end, we aligned the sequences of Nme1Cas9, SmuCas9, and Nme2Cas9 to examine the key interaction interfaces in the PLL and the RuvC-BH linker. We found that, while the sequence of the RuvC-BH linker is highly conserved, the PLL region of Nme2Cas9 is distinct from the two other orthologs. Nme2Cas9 has one less Gly residue and an insertion of three contiguous residues (VKH) compared to Nme1Cas9 and SmuCas9 (Figure 6A). We hypothesized that these sequence differences result in the inability of AcrIIC5 to inhibit Nme2Cas9. To test this hypothesis, we constructed three variants of Nme2Cas9. The first variant has a glycine insertion between residues A841 and H842 to mimic the position of the glycine in Nme1Cas9 (designated as G842ins), the second variant has a deletion of the three additional VKH residues (VKHdel), and the third comprises both G842ins and VKHdel. We found that the single glycine insertion was sufficient for the activity of Nme2Cas9 to be inhibited by AcrIIC5, although its activity was slightly reduced (Figure 6B, Supplementary Figure S6), whereas the VKHdel mutation on its own had no effect. This implies that the PLL, in particular Gly842 of Nme1Cas9, is critical for AcrIIC5 inhibition, which explains why AcrIIC5 can inhibit Nme1Cas9 but not Nme2Cas9.

**Figure 6.**
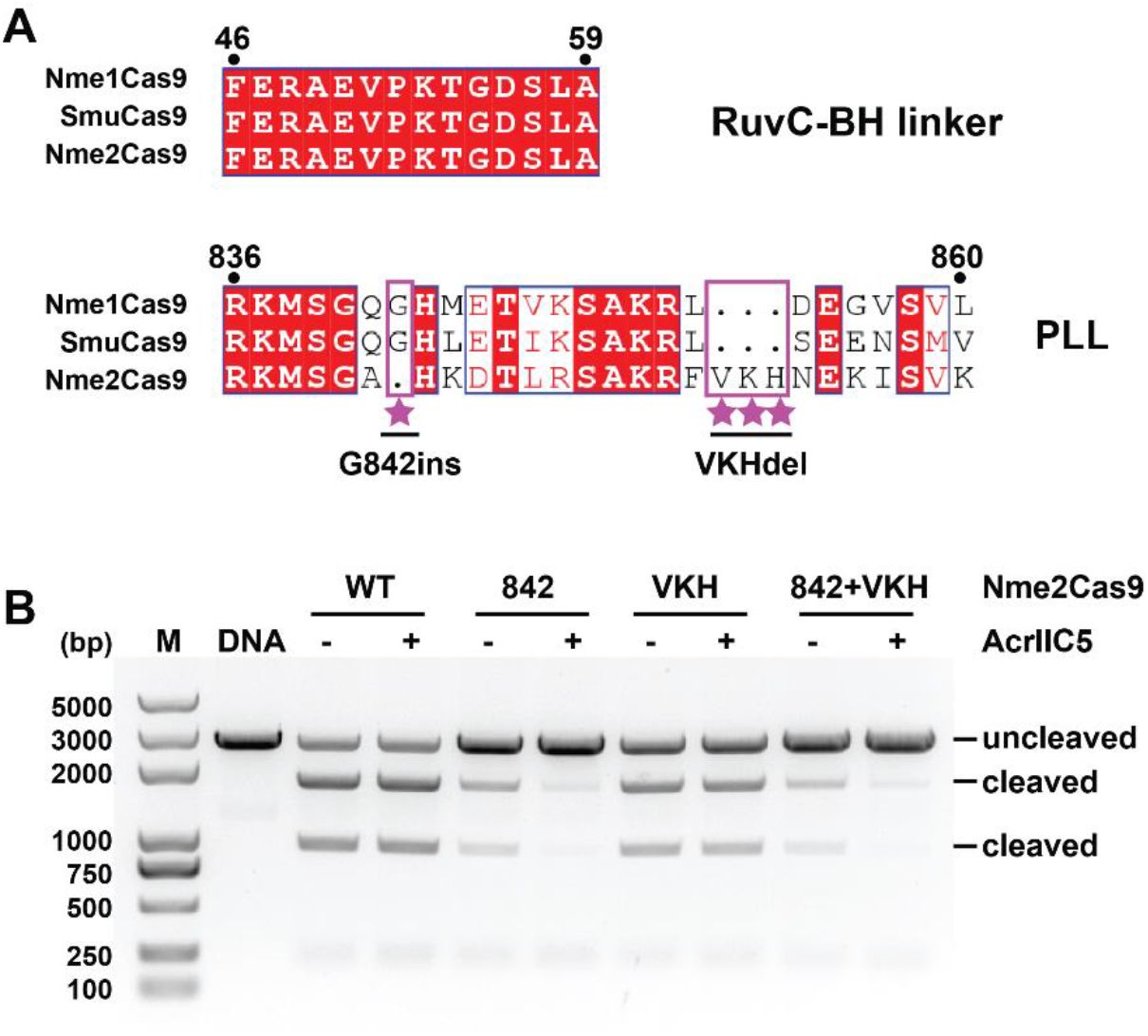
Engineering of Nme2Cas9 based on sequence alignment. (A) Sequence alignments of the RuvC-BH linker and PLL regions of Nme1Cas9, SmuCas9, and Nme2Cas9. Distinct features of Nme2Cas9 are highlighted with violet rectangle and labeled with stars. Three variants of Nme2Cas9, designated as G842ins, VKHdel and (G842ins+VKHdel) double mutants were designed based on this sequence alignment. (B) DNA cleavage activities of Nme2Cas9 variants in absence or presence of AcrIIC5 using a linearized plasmid substrate. Cleavage was performed at 37°C for 10 min. The ability of AcrIIC5 (molar ratio Cas9:Acr of 1:10) to inhibit wild-type (WT), glycine 842 insertion (842), three-residue deletion (VKH), or 842+VKH was measured 1% agarose gel stained with ethidium bromide. The figure is a representative of three replicates.

## Discussion

Our structural and biochemical data reveals that the monomeric AcrIIC5 binds to type II-C Cas9 in the cleft between the WED and PI domains where it mimics the shape and charge of a DNA duplex to occupy the DNA binding pocket and conceal the PAM recognition site. Thus, AcrIIC5 inactivates Nme1Cas9 by blocking the DNA binding. Importantly, the PLL and the RuvC-BH linker of Cas9 – the two major binding interfaces of AcrIIC5 – are both essential for the target dsDNA unwinding (47-50). Upon target DNA binding, residues E845 and T846 within the PLL are responsible for stabilizing the phosphate group of the first nucleotide in the unwound target strand (TS) (Supplementary Figure S4). Residue K53 within the RuvC-BH linker inserts between the first and the second nucleotides of the unwound non-target strand (NTS), preventing the re-annealing of TS and NTS. Thus, K53 facilitates positioning of NTS into the RuvC active pocket (Supplementary Figure S4). However, in the presence of AcrIIC5, these crucial amino acids interact with the residues D36 and E37 of AcrIIC5, and are therefore not able to participate in the early stages of target binding. Altogether, our results show that AcrIIC5 acts as a dsDNA mimic with a novel protein fold, preventing dsDNA recognition and the melting of double-stranded target DNA by type II-C Cas9 surveillance complexes. These characteristics make AcrIIC5 a potent type II-C anti-CRISPR protein.

DNA mimicry is a common strategy used by bacteriophage proteins, including previously characterized anti-CRISPRs. For example, AcrF2 is a dsDNA mimic that prevents target recognition in the type I-F CRISPR-Cas system (51,52), and both AcrIIA2 and AcrIIA4 inhibit type II-A Cas9 by blocking target dsDNA recognition (23,44-46). Our new work presents the first example of a type II-C Acr that mimics DNA to block the target DNA engagement of type II-C Cas9. One common property of DNA-mimic Acrs is that they are acidic proteins with large negatively-charged surfaces, resembling the shape and charge distribution of DNA (Supplementary Figure S7). In addition, these Acrs only bind to sgRNA-bound Cas9, probably because the DNA binding surface on Cas9 only forms after sgRNA loading. DNA mimics directly compete for the DNA binding site, in particular the PAM recognition site, thus rendering the surveillance complex unable to bind or cleave target nucleic acids.

AcrIIC5 from *Simonsiella muelleri* inhibits both Nme1Cas9 and SmuCas9, with the inhibition of SmuCas9 being more potent (Figure 1A) – perhaps unsurprisingly, since this ortholog is native to the same species from which the Acr was isolated and thus likely represents its natural target. The double mutant D36A/E37A abolished AcrIIC5’s ability to inhibit Nme1Cas9, but only modestly reduced its inhibition of SmuCas9 (Figure 4A, Supplementary Figure S5), suggesting that other as-yet unidentified residues also contribute to the interaction between AcrIIC5 and SmuCas9. The sequences of SmuCas9 and Nme1Cas9 are highly similar (74% similarity), in particular the PLL and RuvC-BH linker (Figure 6A), implying that AcrIIC5 interacts with SmuCas9 in a similar manner, but more strongly than Nme1Cas9. The ability of AcrIIC5 to inhibit Nme1Cas9 but not Nme2Cas9 was interesting to us, since these two paralogs share 89% similarity. We were able to pinpoint this striking difference to a single glycine residue in the PLL that is present in Nme1Cas9 but absent in Nme2Cas9, which raises the possibility that Cas proteins may evolve to escape Acr binding. In addition, our elucidation of the inhibition mechanism of AcrIIC5 may facilitate the development of new tools to regulate genome editing or better understand type II-C Cas9 mechanisms of action.

## Data Availability

The maps used to generate the cryo-EM volume have been deposited in the Electron Microscopy Data Bank (EMDB) with accession number EMD-34832. Real-space refined coordinate for Nme1Cas9-sgRNA-AcrIIC5 complex has been deposited in the Protein Data Bank (PDB) with accession number 8HJ4.

## Supplementary Data

Supplementary Data are available at NAR online.

## Acknowledgement

We are grateful to the staff of the Center for Biological Imaging, Core Facilities for Protein Science at the Institute of Biophysics, Chinese Academy of Sciences for support to collect cryo-EM data. We thank Dr. April Pawluk in Life Science Editors for help with editing the manuscript.

