## Supplements for "Anti-CRISPR AcrIIC5 is a dsDNA mimic that inhibits type II-C Cas9 effectors by blocking PAM recognition"

**Supplementary Table S1. RNA and DNA sequences used in this paper**

| DNA IDENTIFIER | Sequence (5' to 3') | SOURCE |
|---|---|---|
| sgRNA of Nme1Cas9, Nme2Cas9, SmuCas9 | GGUCACUCUGCUAUUUAACUUUACGUUGUAGCUCCCUUUCUCGAAAGAGAACCGUUGCUACAAUAAGGCCGUCUGAAAAGAUGUGCCGCAACGCUCUGCCCCUUAAAGCUCCUGCUUUAAGGGGCAUCGUUUAUC | Sangon Biotech |
| Nme1Cas9-sgRNA-DNA (partially duplexed) | TS: TAAAATCATATGTGTGATTAAATGTTAGAGTGACC | Sangon Biotech |
| | NTS: ATATGATTTTA | Sangon Biotech |
| Nme1Cas9-sgRNA-DNA (linear plasmid target) | TS: TAAAATCATATGTGTGATTAAATGTTAGAGTGACC | Sangon Biotech |
| | NTS: GGTCACTCTAACATTTAATCACACATATGATTTTA | Sangon Biotech |
| Nme1Cas9-sgRNA-DNA (Cy3-labeled dsDNA for EMSA) | TS: Cy3-CTCAGTGATCTAAAATCATATGTAAAGTTAAATAGCAGAGTGACCTGTCATGA | Sangon Biotech |
| | NTS: TCATGACAGGTCACTCTGCTATTTAACTTTACATATGATTTTAGATCACTGAG | Sangon Biotech |
| Nme2Cas9/SmuCas9-sgRNA-DNA (linear plasmid target) | TS: TAACTGGGCCTGTAAAGTTAAATAGCAGAGTGACC | Sangon Biotech |
| | NTS: GGTCACTCTGCTATTTAACTTTACAGGCCCAGTTA | Sangon Biotech |

**Supplementary Table S2. Cryo-EM data collection, refinement and validation statistics**

| | Nme1Cas9-sgRNA-AcrIIC5 |
|---|---|
| **Data collection and processing** | |
| Magnification | 130,000 |
| Voltage (kV) | 300 |
| Electron exposure (e-/Å$^2$) | 60 |
| Defocus range (μm) | -1.0 to -1.6 |
| Pixel size (Å) | 1.04 |
| Symmetry imposed | C1 |
| Initial particle images (no.) | 2,685,237 |
| Final particle images (no.) | 291,265 |
| Map resolution (Å) | 3.09 |
| FSC threshold | 0.143 |
| Map resolution range (Å) | 3 to 8 |
| **Refinement** | |
| Initial model used (PDB code) | 6JDQ |
| Model resolution (Å) | 3.4 |
| FSC threshold | 0.5 |
| Map sharpening B factor (Å$^2$) | -57.4 |
| Model composition | |
| Non-hydrogen atoms | 11,652 |
| Protein residues | 1166 |
| Nucleotides | 113 |
| Ligands | 0 |
| Mean B factors (Å$^2$) | |
| Protein | 66.49 |
| Nucleotides | 97.32 |
| R.m.s. deviations | |
| Bond lengths (Å) | 0.003 |
| Bond angles (°) | 0.600 |
| Validation | |
| MolProbity score | 2.07 |
| Clashscore | 12.75 |
| Poor rotamers (%) | 0.00 |
| Ramachandron Plot | |
| Favored (%) | 92.89 |
| Allowed (%) | 6.85 |
| Outliers (%) | 0.26 |

### Supplementary Figures

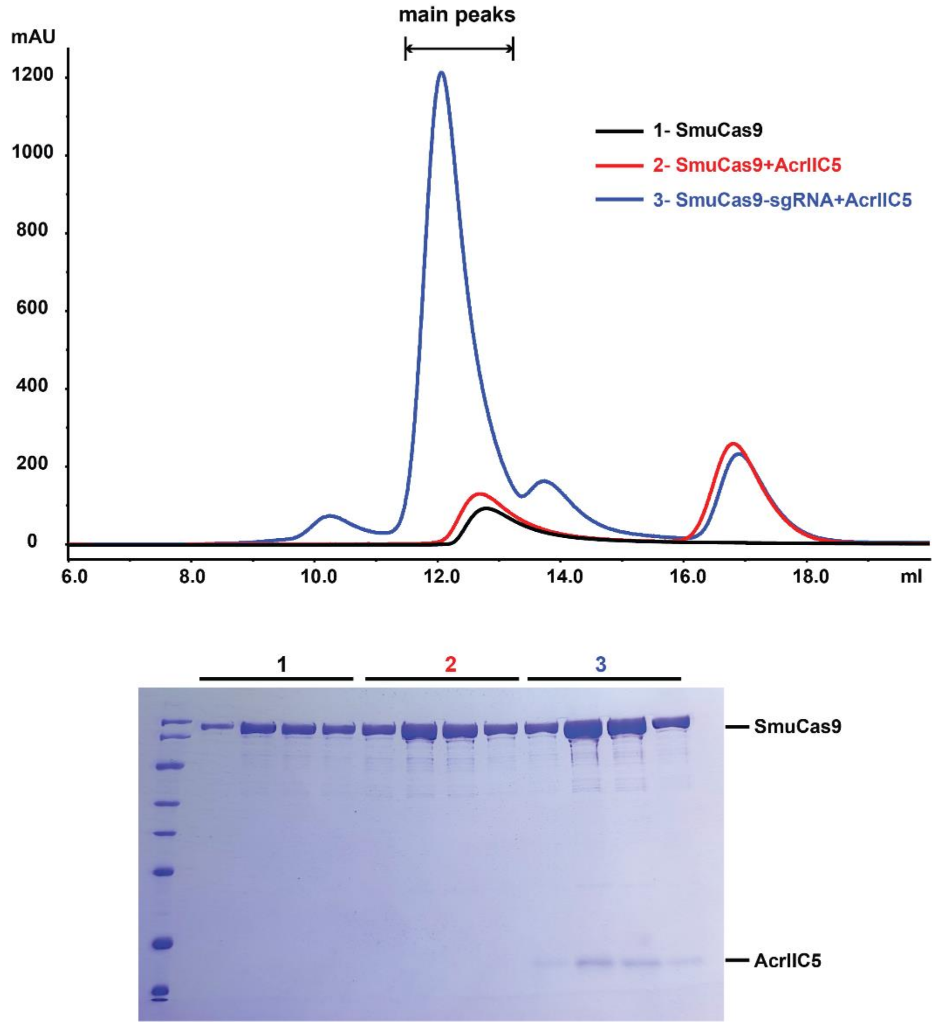


**Supplementary Figure S1. AcrIIC5 binds to sgRNA-bound SmuCas9 but not apo-SmuCas9**

Size-exclusion chromatography curves of apo-SmuCas9 (black), SmuCas9+AcrIIC5 (red), and SmuCas9-sgRNA+AcrIIC5 (blue) are overlaid. Fractions from the Cas9-containing main peaks were characterized by SDS-PAGE followed by Coomassie blue staining.

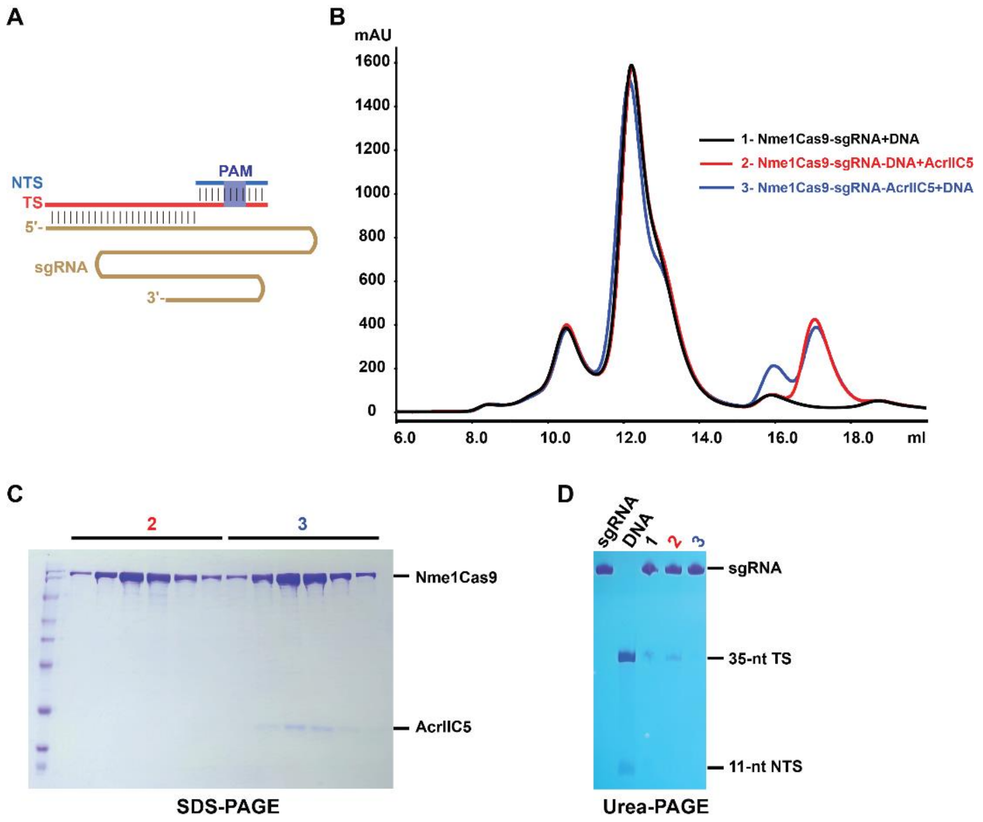


**Supplementary Figure S2. AcrIIC5 competes dsDNA for binding to Nme1Cas9-sgRNA**

(A) A schematic of the sgRNA and partially duplexed dsDNA used for this assay.

(B) Size-exclusion chromatography curves of Nme1Cas9-sgRNA+DNA (black), Nme1Cas9-sgRNA-DNA+AcrIIC5 (red) and Nme1Cas9-sgRNA-AcrIIC5+DNA (blue) were overlaid.

(C) Fractions from the main peaks of the SEC shown in (B) were characterized by SDS-PAGE and stained with Coomassie blue. The colors and numbers of labels on the top of the gel are consistent with the ones of the SEC traces shown in (B).

(D) Samples extracted from fractions of the main peaks were characterized by TBE-Urea-PAGE and stained with toluidine blue to visualize the nucleic acids present in each peak fraction. The sample numbers and colors of the labels are consistent with the ones of the SEC curves from (B).

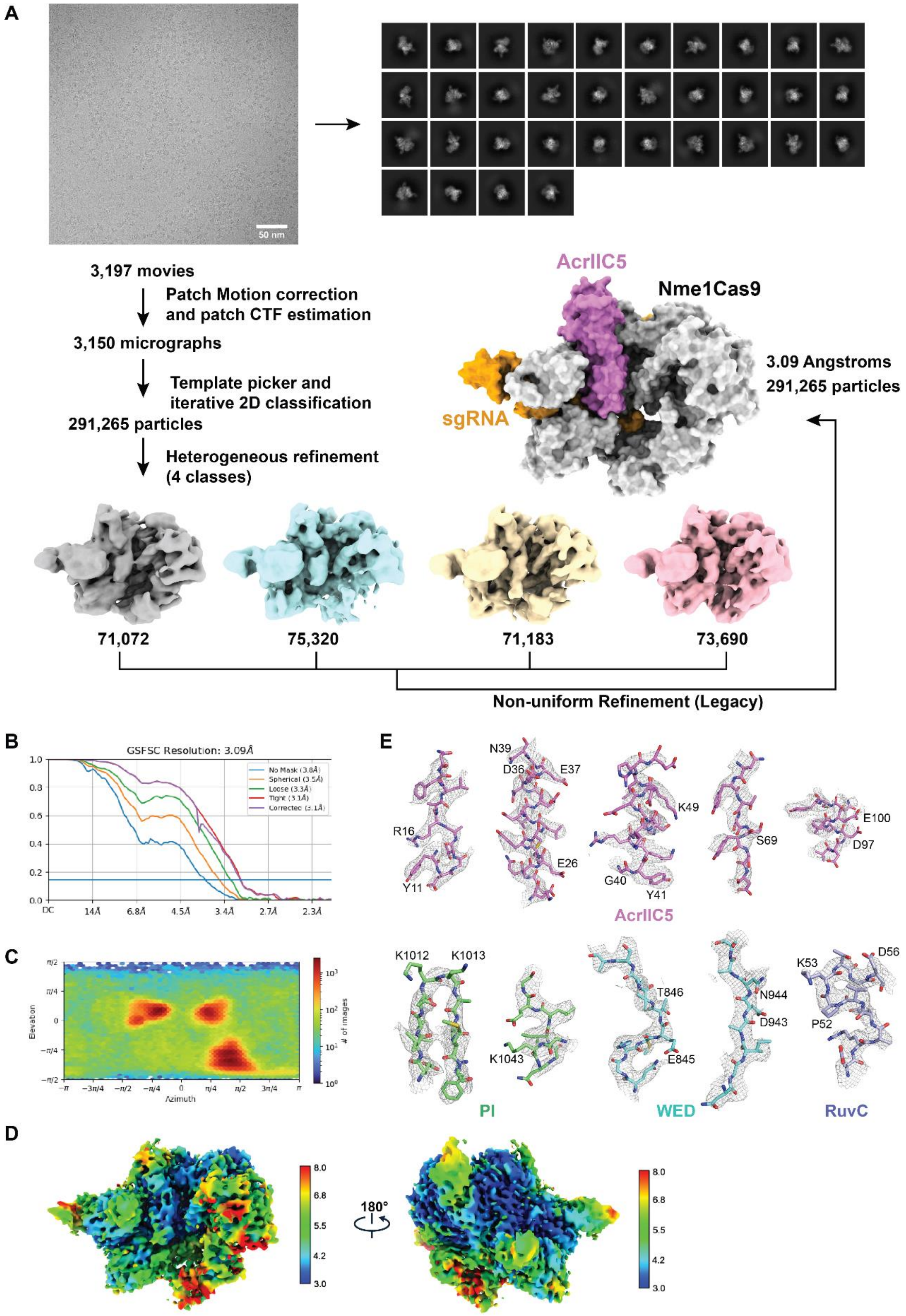


**Supplementary Figure S3. Cryo-EM single particle reconstruction of Nme1Cas9-sgRNA-AcrIIC5 complex**

(A) Workflow of the cryo-EM image processing and 3D reconstruction for the Nme1Cas9-sgRNA-AcrIIC5 complex. The final electron density maps of different chains are colored separately.

(B) Fourier Shell Correlations (FSC) of Nme1Cas9-sgRNA-AcrIIC5 complex reconstruction, with the gold-standard cutoff (FSC=0.143) marked with a horizontal blue line.

(C) Direction distribution plot.

(D) Final electron density map showing local resolution. Two opposite views are displayed.

(E) Cryo-EM densities of residues involved in Cas9:AcrIIC5 interactions.

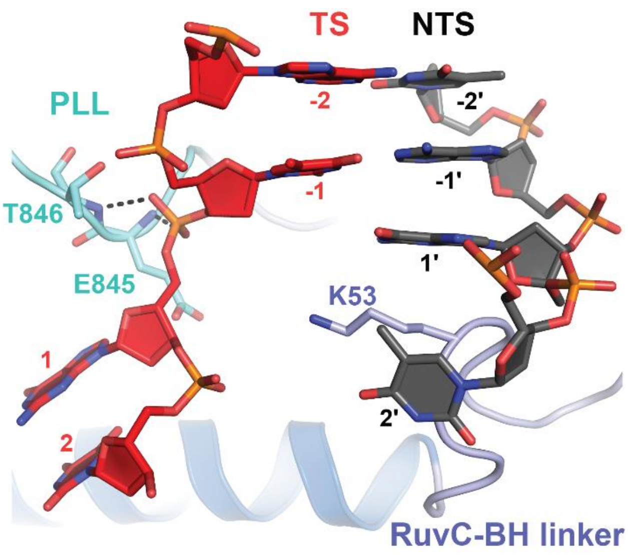


Nme1Cas9-sgRNA-AcrIIC3-dsDNA

**Supplementary Figure S4. The PLL and RuvC-BH linker are critical for dsDNA unwinding**

In the structure of Nme1Cas9 H588A-sgRNA-AcrIIC3-dsDNA (PDB: 6JE4), the hydrogen bonds between residues from the PLL and the TS are shown as black dashed lines. Residue K53 from the RuvC-BH linker, which inserts between the first and second bases from the NTS, is shown as stick.

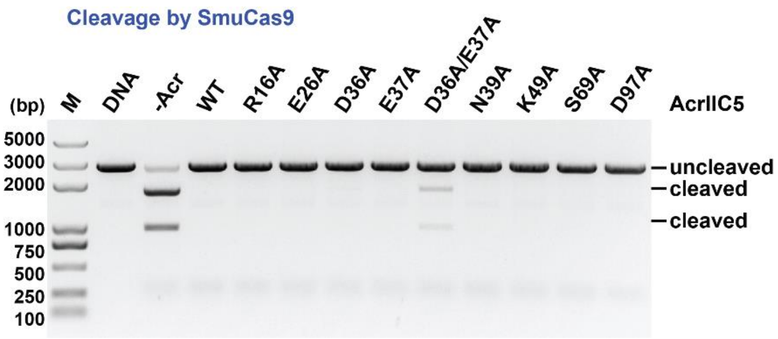


**Supplementary Figure S5. Cleavage assay of SmuCas9 inhibited by single or double mutants of AcrIIC5 using linearized plasmid substrate**

In vitro DNA cleavage assay of SmuCas9 using a linearized plasmid substrate in the absence or presence of wild-type or indicated mutants of AcrIIC5. The molar ratio of Cas9 to AcrIIC5 is 1:10. Cleavage was performed at 37°C for 60 min. The figure is a representative of three replicates.

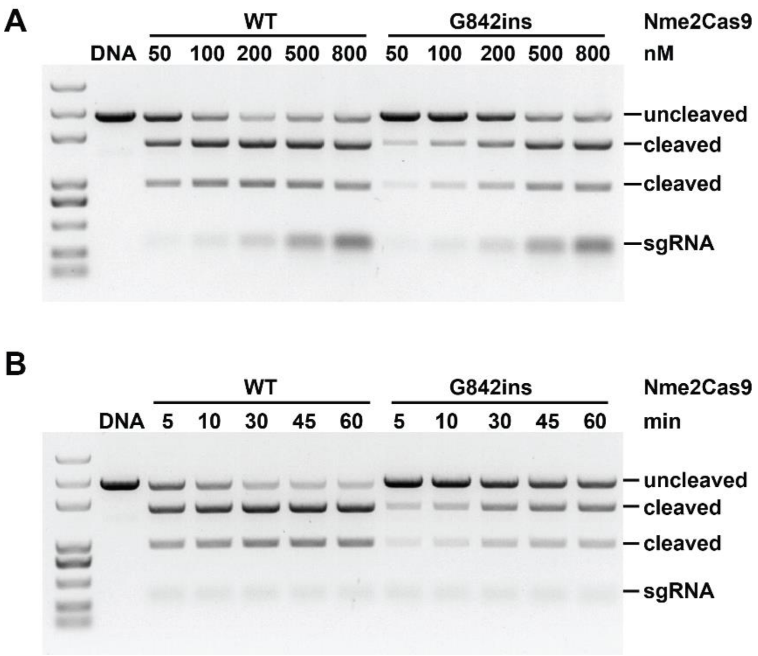


**Supplementary Figure S6. Comparison of the activities of WT and G842ins mutant of Nme2Cas9.**

(A) In vitro cleavage of linearized plasmid substrate with WT and mutant of Nme2Cas9 in an increase gradient.

(B) Time-course cleavage assay of WT and mutant of Nme2Cas9 using linearized plasmid substrate.

The figures are representatives of three replicates.

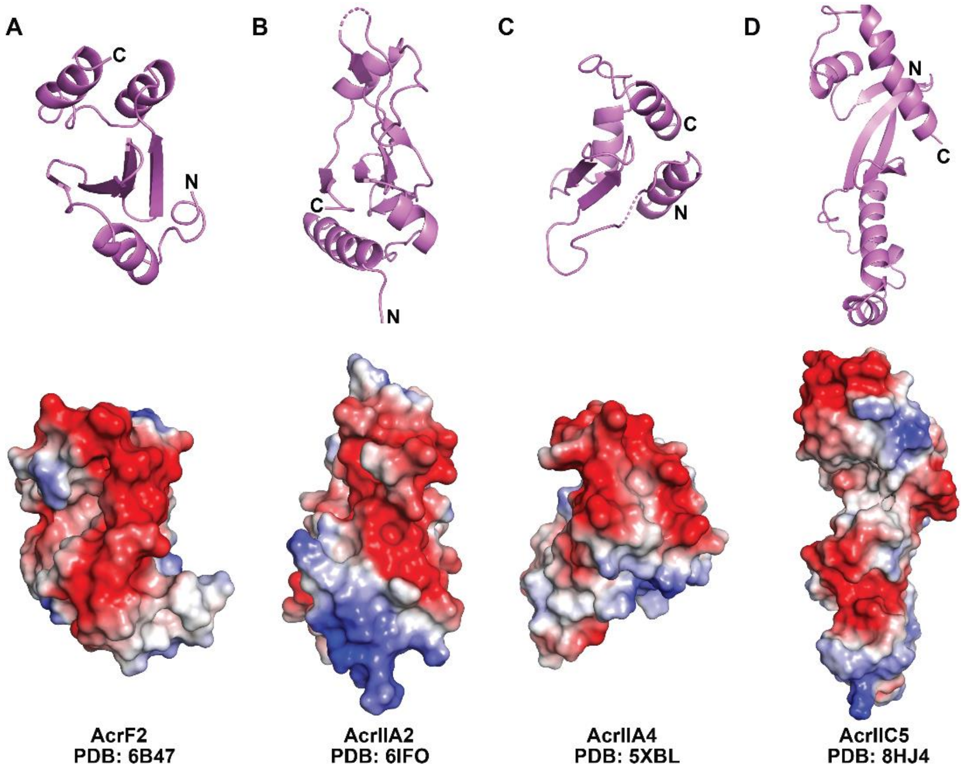


**Supplementary Figure S7. Comparison of the structures and charge distributions of various PAM-mimic anti-CRISPR proteins**

Anti-CRISPR proteins shown as cartoon are on the top and their electrostatic surface potentials are displayed at the bottom with the positive charge colored in blue and negative charge in red.
